# Comparative analysis of whole genome sequences of *Leptospira* spp. from RefSeq database provide interspecific divergence and repertoire of virulence factors

**DOI:** 10.1101/2021.01.12.426470

**Authors:** Mohd Abdullah, Mohammad Kadivella, Rolee Sharma, Mirza. S. Baig, Syed M. Faisal, Sarwar Azam

**Affiliations:** Computational Biology Lab, National Institute of Animal Biotechnology, Hyderabad, India; Laboratory of Vaccine Immunology, National Institute of Animal Biotechnology, Hyderabad, India; Department of Biosciences, Integral University, Lucknow, India; Centre for Biosciences and Biomedical Engineering, Indian Institute of Technology, Indore, MP, India; Department of Biomedical Engineering, Indian Institute of Technology, Hyderabad, India

**Keywords:** Comparative genomics, *Leptospira*, virulence factors

## Abstract

Leptospirosis is an emerging zoonotic and neglected disease across the world causing huge loss of life and economy. The disease is caused by *Leptospira* of which 605 sequenced genomes representing 72 species are available in RefSeq database. A comparative genomics approach based on Average Amino acid Identity (AAI), Average Nucleotide Identity (ANI), and Insilco DNA-DNA hybridization provide insight that taxonomic and evolutionary position of few genomes needs to be changed and reclassified. Clustering on the basis of AAI of core and pan-genome contradict clustering pattern on basis of ANI into 4 clusters. Amino acid identity based hierarchical clustering clearly established 3 clusters of *Leptospira* correlating with level of virulence. Whole genome tree supported three cluster classifications and grouped *Leptospira* into three clades termed as pathogenic, intermediate and saprophytic. *Leptospira* genus consist of diverse species and exist in heterogeneous environment, it contains relatively large and closed core genome of 1038 genes. Analysis provided pan genome remains open with 20822 genes. COG analysis revealed that mobilome related genes were found mainly in pan-genome of pathogenic clade. Clade specific genes mined in the study can be used as marker for determining clade and associating level of virulence of any new *Leptospira* species. Many known *Leptospira* virulent genes were absent in set of 78 virulent factors mined using Virulence Factor database. A deep search approach provided a repertoire of 496 virulent genes in pan-genome. Further validation of virulent genes will help in accurately targeting pathogenic *Leptospira* and controlling leptospirosis.

**Graphical Abstract:** 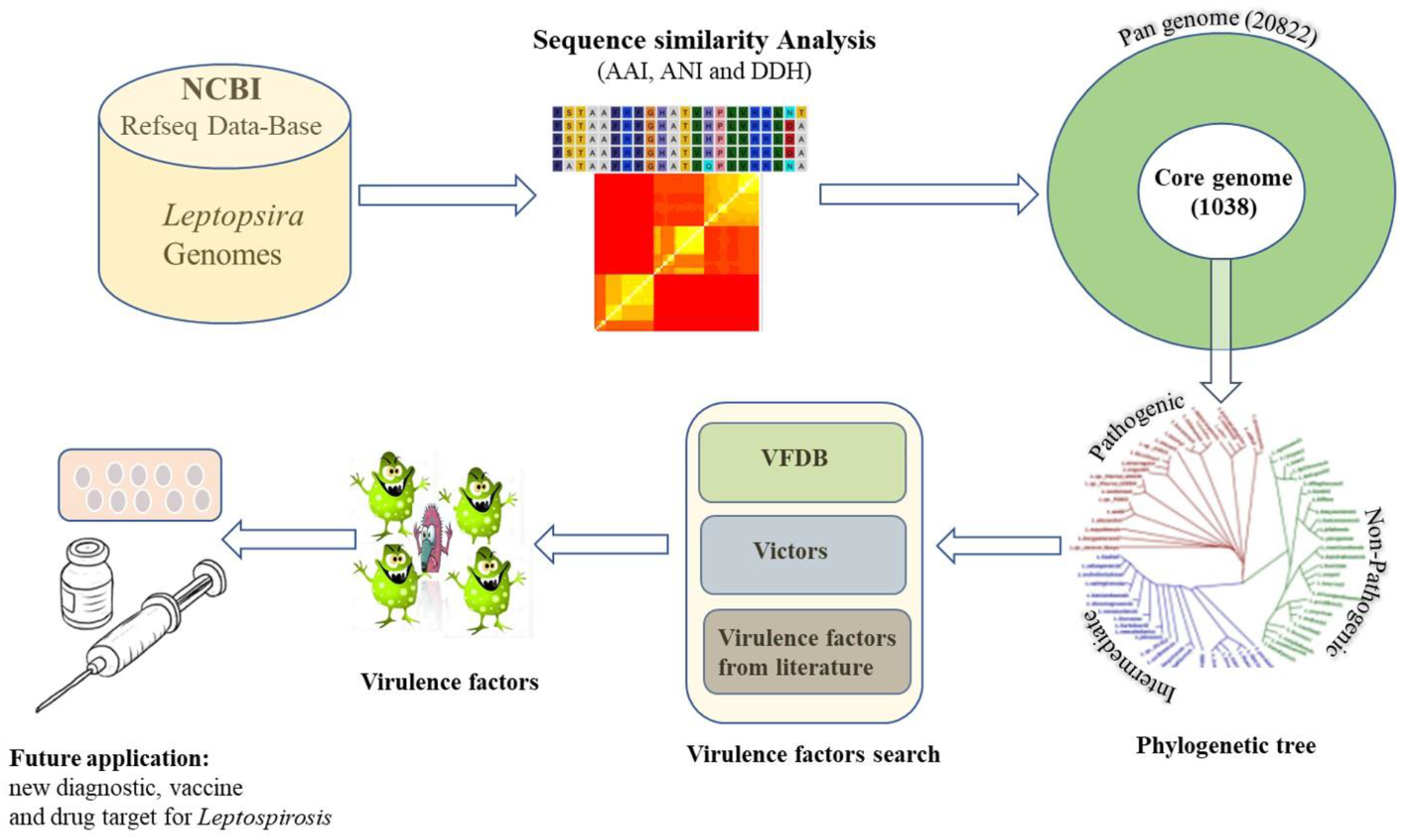

## 1. INTRODUCTION

With the reduction in the cost of Next generation sequencing (NGS) technologies, it has become most suitable tool for dissecting the complete genome of species. Recently, NGS has been applied in monitoring and characterizing the outbreak strain of bacteria as well as viruses. Complete genome sequences help in establishing, verifying and updating knowledge of known taxonomic positions, evolutionary relationships, sub-typing and another genomic features. It also provides supporting information on virulence factors, pathogenesis, metabolism, host-pathogen interaction, host-environment interaction etc. Core and pan-genome at the level of species and genus provide conservation, selection and addition of genes in the genome(Chen et al., 2018; Grote et al., 2012; Guimaraes et al., 2015). Identifying unique genes may define phenotypic feature of the species(Bochner, 2009). However, genes common and exclusive to a group of species provide feature of the whole group. This group or clad specific genes are normally termed as gene gain and can be developed as marker for characterizing the species(Li et al., 2017).

Leptospirosis is a worldwide-distributed zoonotic and infectious diseases(Faisal et al., 2016). It is one of the neglected and emerging disease of public health status with respect to morbidity and mortality both in animals and humans (Karpagam and Ganesh, 2020). Only in human, average morbidity of 1.03 millions per year has been reported which causes around sixty thousands deaths across the world(Karpagam and Ganesh, 2020). The disease is caused by gram negative bacteria called *Leptospira*. *Leptospira* are known to have 3 groups i.e. Pathogenic, Intermediate and saprophytic based on their level of virulence. Intermediate species shows moderate pathogenicity in both human and animals. Various serological assays have been developed to test the pathogenicity and classify the *Leptospira* in groups(Cerqueira and Picardeau, 2009). However, serological groups or serotype does not clearly define species. So, 16S RNA sequencing or Multi Locus Sequence Typing (MLST) were used initially to differentiate at species level(Ahmed et al., 2006; Azam et al., 2016). First genome of *Leptospira* was sequenced and characterized using first generation sequencing method in 2003 (Ren et al., 2003). Later, due to low cost NGS platforms, many *Leptospira* genomes have been sequenced across the world in recent years. With availability of genomes, many comparative genomics studies were carried out to understand pathogenicity and evolution of Leptospira (Bulach et al., 2006; Fouts et al., 2016; Lehmann et al., 2013; Nascimento et al., 2004; Zhong et al., 2011). Comparative genomics also revealed that pathogenic Leptospira may undergo genome reduction(Bulach et al., 2006). Sequencing of saprophytic species and its comparison provided specific genes for pathogenic clades (Fouts et al., 2016; Picardeau et al., 2008; Xu et al., 2017). In another study, Xu et al. reported genome sequence of 102 isolates and established Core and pan genome with18 species of Leptospira(Xu et al., 2016). It also suggested that pathogenic Leptospira acquires virulence factors through horizontal gene transfer. Recently, Vincent et. Al., sequenced 30 novel species and revisited the taxonomy and evolution of pathogenicity of Leptospira (Vincent et al., 2019).

At present ~ 600 genomes which represent more than 70 species of *Leptospira* have been sequenced and deposited in NCBI((ftp://ftp.ncbi.nlm.nih.gov/genomes/refseq/bacteria/). This provides an opportunity to revisit and characterize the *Leptospira* genus comprehensively and compare the genomic features of all available species. In this study, whole genome sequences were used to calculate genome to genome distance, nucleotide similarity, core and pan genome of whole genus for a better understanding about this pathogen. The data were also analysed to extract specific genes for marker development and identify virulence factors which can used for targeting leptospirosis through development of novel vaccine and diagnostic.

## 2. Material and methods

### 2.1 Data collection and genome sequences of Leptospira species

All the latest assemblies and annotations of representative genome of each species of *Leptospira* were downloaded from NCBI Reference Sequence (RefSeq) Database (ftp://ftp.ncbi.nlm.nih.gov/genomes/refseq/bacteria/). Each species with its gene content and other features like GC content, length of genome assembly, host and place from which it was isolated, etc. were tabulated. Unusual genome or outlier on basis of gene content was also searched using bubble sort method.

### 2.2 Average Nucleotide Identity (ANI), Average Amino Acid Identity (AAI) and Insilco DNA-DNA hybridisation between *Leptospira* genomes

Average Nucleotide Identity (ANI) between representative genomes of each species of *Leptospira* were calculated through PYANI V0.2.10 software (https://github.com/widdowquinn/pyani.git). ANI also called ANIb as blast software were used in the analysis. ANI square matrix generated through Pyani were clustered and heatmap was generated using scripts of GET_HOMOLOGUES(Contreras-Moreira and Vinuesa, 2013) software (get_homologus-x86_64) and R (V.3.6.2)(Pritchard et al., 2019). Average Amino Acid Identity (AAI) between species were calculated on basis of homologous genes using scripts of GET_HOMOLOGUES(Contreras-Moreira and Vinuesa, 2013). In-silico DNA-DNA hybridisation (DDH) were calculated using online version of Genome-to-Genome Distance Calculator(GGDC v2.1, https://ggdc.dsmz.de/ggdc.php#)(Meier-Kolthoff et al., 2013; Meier-Kolthoff et al., 2014).

### 2.3 Clustering and analyses of core and pan genomes

Core and pan genome of *Leptospira* genus was determined using GET_HOMOLOGUES package (Contreras-Moreira and Vinuesa, 2013). Orthologous genes across all the representative species was obtained with Bidirectional best hits (BDBH) using BLASTP of NCBI-BLAST+ vs2.10.0 (ftp://ftp.ncbi.nlm.nih.gov/blast/executables/blast+/LATEST/). Script “get_homologues.pl” with parameter −C −70, −E 1e-05, −F 1.5, −M (for OMCL), −m local and −t 0 were used to cluster the genes. Tetteline and Willin brook fits were used to estimate core genome size and Tettiline formulae was applied to estimate pan-genome size using script “plot_pancore_matrix.pl”. A shell script “hcluster_pangenome_matrix.sh” was used for clustering of species and generation of heatmap. Clad specific genes were identified using script “parse_pangenome_matrix.pl” with setting −m that is taking pan-genome matrix as input file, −g finding unique genes present in −A which is list of species in a clad, but not in −B which is list of species in another clad. Sequences of clad specific genes were extracted using in-house perl script.

### 2.4 Phylogenomic analysis of *Leptospira*

Single copy orthologs of *Leptospira* were obtained from core genome. MAFFT(Katoh and Standley, 2014) v79 was used for multiple sequence alignment (MSA) of each orthologous group which were further processed and concatenated using G-block(Talavera and Castresana, 2007). The alignment file converted to Phylip format using script “fasta2phylip.pl”. RAxML(Stamatakis, 2014) was run on alignment for a rapid Bootstrap analysis and search for the best-scoring ML tree with PROTGAMMALG. The output of RAxML was visualized using Dendroscope(Huson et al., 2007). The whole analysis was repeated after including *Leptonema illini* as outgroup.

### 2.5 Cluster of Orthologous Groups (COGs) analysis

Different proteome dataset of core, soft core and pan-genome of each clad of *Leptospira* were subjected to blast against COGs profiles. COGs profile was obtained from the conserved domain database (CDD) (ftp://ftp.ncbi.nih.gov/pub/mmdb/cdd) (Tatusov et al., 2000). RPS-BLAST+ v2.10.0 was used for similarity search with an e-value cut-of 0.01. Protein sequence having at least 25% identity and 70% alignment length with corresponding COG Profile were assigned to the functional group.

### 2.6 Pathway analysis using GhostKOALA

Gene’s association with pathway were searched using KEGG database (https://www.kegg.jp/ghostkoala/). Ghostkoala annotation server with option “genus_prokaryote + family eukaryote” was used to perform KO analysis to assign individual gene functions. Genes were annotated and associated with its pathways using BRITE, pathway and module reconstruction tool of KEGG MAPPER.

### 2.7 Virulence factor analysis

Genes were searched for orthologs present in core dataset of Virulence factor database (VFDB) (http://www.mgc.ac.cn/VFs/download.htm)(Chen et al., 2005) using BLAST+ v2.10.0. Stringent criteria of 40% identity and 70% alignment coverage were used to establish the orthology. Similar analysis was repeated with Victors database (http://www.phidias.us/victors/download.php)(Sayers et al., 2019). Further, virulence factors of *Leptospira* from different published literature were manually compiled, curated and their sequences were fetched from Uniprot (https://www.uniprot.org/) and NCBI (https://www.ncbi.nlm.nih.gov/). Pangenome of different clades were searched against these known genes.

## 3. Results

### 3.1 Leptospira species are extensively sequenced

As per NCBI database, a total of 624 genomes representing 73 species of *Leptospira* genus were sequenced and available (**Supplementary Table S1**). Maximum number of the sequenced genomes belongs to pathogenic species such as *Leptospira interrogans* (305) and *Leptospira borgpetersonii* (44). Non-pathogenic *Leptospira* species from soil, water and environment are poorly represented and only few genomes are sequenced. Most of the sequenced genomes are isolated from human host followed by Cattle. We analysed all the representative genomes of 73 species available at NCBI. On an average *Leptospira* species contain ~3800 genes. However, *Leptospira borgpetersoni* contain minimum number of genes and *L. macculloughii* contain highest number of genes (6390). A bubble sort analysis clearly revealed that *L. macculloughii* is the only outlier species, hence, it was excluded from downstream analysis (**Supplementary Figure S1**). Further we observed *Leptospira* genomes are AT rich, where GC content vary from 35.02 - 47.77%.

### 3.2 Assessment of core and pan genomes

To assess the diversity in gene content of *Leptospira* genus, core and pan genome were calculated. The pan-genome of *Leptospira* never saturated and increased with 91 genes, even after the addition 72^nd^ species. Therefore, pan-genome of *Leptospira* remain open (**Figure 1b**). However, core genome decreased drastically with addition of initial few genomes and started stabilizing after addition of 51th genome. As per Tettelin formulae(Tettelin et al., 2008), graph become parallel to X axis (**Figure 1a),** indicating that core genome is closed one. In this study, Pan-genome and core genome consist of 20822 and 1038 genes respectively. Accessory genome consists of 6488 genes (31.1%) in the shell and 12486 genes (59.9%) in cloud (**Figure1c, d**).

**Figure 1:**
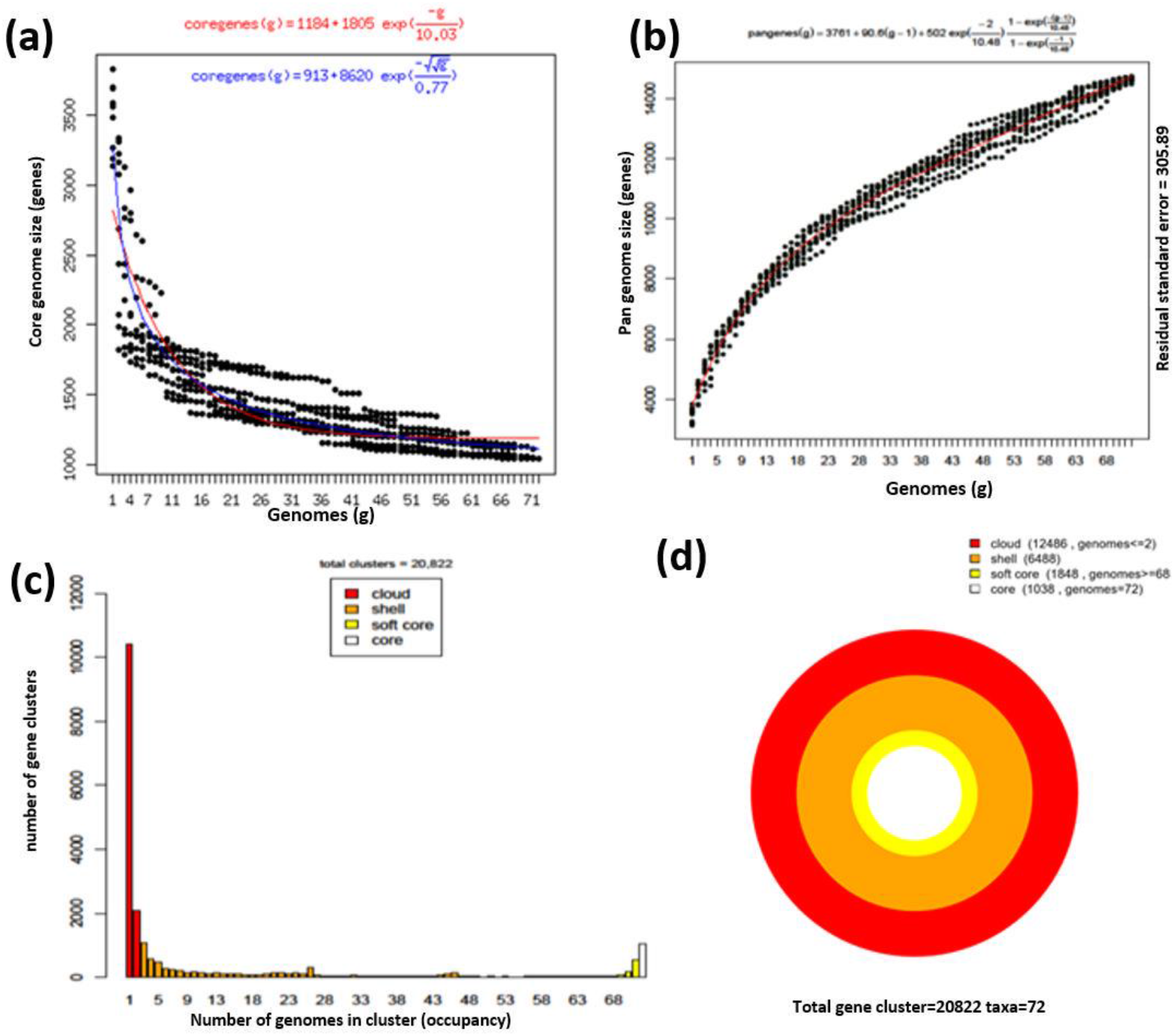
Comparative genomic analysis of Leptospira species. **(a)** Core genome estimation with Tettelin (blue) and Willienbrock (red) fits **(b) Evolution** of Pan genome size with the Tettelin fit (red)**. (c) Bar plot of pan genome:** red bars represent cloud genes, orange bars represent shell genome, yellow bars represent softcore and white bar represents core genome. **(d) Circular plot of pan genome:** innermost circle (in white) represents core genome. The 2^nd^ innermost circle (in yellow) represents gene in softcore whereas 2^nd^ outermost circle (in orange) represent shell and the outermost circle (in red) represent cloud genes which are unique or present in maximum of two genomes.

### 3.3 Similarity and distances between *Leptospira* genomes

#### ANIb analysis across Leptospira

After excluding *L. macculloughii*, remaining 72 representative genomes of *Leptospira* were compared with each other for genomic similarity. ANIb values summarized in table (**Supplementary Table S2**) shows least nucleotide identity (68.92%) exist between *Leptospira ryugenii* and *Leptospira santrosi*. Heatmap (**Supplementary Figure S2**) generated using ANI value clearly depicts 4 optimal groups in dendrogram supported with silhouette width calculation (**Supplementary Figure S3b**). Group (A) and (B) consist of 24 and 21 species respectively. Group (C) consisting of 13 members with at least 80% identity to each other whereas group (D) with 14 members showed very less identity (69-73%). Group A, B, and C are visible in heatmap with yellow to red colour. However, 8 pairs of genomes showed more than 95% average nucleotide identity and visible as light yellow to red in heatmap like other self-compared genomes showing 100% nucleotide identity. *L*. sp. Fiocruz LV4135, *L*. sp. Fiocruz LV3954, and *L*. santarosai showed more than 98% nucleotide identity with each other. Others pair of genome showing more than 95% are *L*. putramalaysiae vs *L*. stimsonii, *L*. weilli vs *L*. sp. P2653, *L*. yasudae vs *L*. dzianensis, *L*. sp. Serovar Kenya vs *L*. borgpetersenii and *L*. kirschneri Vs *L*. sp ZV016.

### 3.4 DDH analysis across Leptospira

In-silico DNA-DNA hybridization (DDH) value with cut-off of 70% are considered for grouping two Leptospira as species. We found 5 such pairs of genomes of which *L*. sp. Fiocruz LV4135, *L*. sp. Fiocruz LV3954 and *L*. santarosai showed very high value of DNA-DNA hybridization with each other **(Supplementary Table S3)**. Other two pair which showed more than 70% hybridization are *Leptospira* sp. Serovar Kenya vs *Leptospira* borgpetersenii(80.1%) and *L*.kirschneri Vs *L*.sp ZV016 (85.7%). The three pair of species i.e. *L*.putramalaysiae vs *L*.stimsonii, *L*.weilli vs *L*.sp. P2653 and *L*.yasudae vs *L*.dzianensis which has very high value of ANI (>95%), showed 63.4%, 62.8%, and 64.9% DDH respectively.

### 3.5 AAI analysis across Leptospira

Average amino acid identity (AAI) was calculated on core genome as well as on pan genome. A core genome based hierarchical clustering revealed 3 optimal clusters (**Figure 2**). Similar clustering was also observed on pan-genome as well (**Supplementary Figure S4**). Three optimal clusters in each analysis were supported with Silhouette width calculation (**Supplementary Figure S3a)**. Each clade was designated on the basis of nature of clustering species. The largest clad was non-pathogenic (saprophytic), clustered with 26 species followed by pathogenic and intermediate consisting 24 and 22 species respectively. On comparison with ANIb dendogram and heatmap **(Supplementary Figure S2**), pathogenic and non-pathogenic clad corresponds to group A, B respectively. Whereas intermediate clad was represented in both group C and D except five species of group D which clustered to non-pathogenic clade. These five species are *Leptospira ryugenii*, *Leptospira ognonensis*, *Leptospira idonii*, *Leptospira ilyithenensis* and *Leptospira kobayashii*.

**Figure 2:**
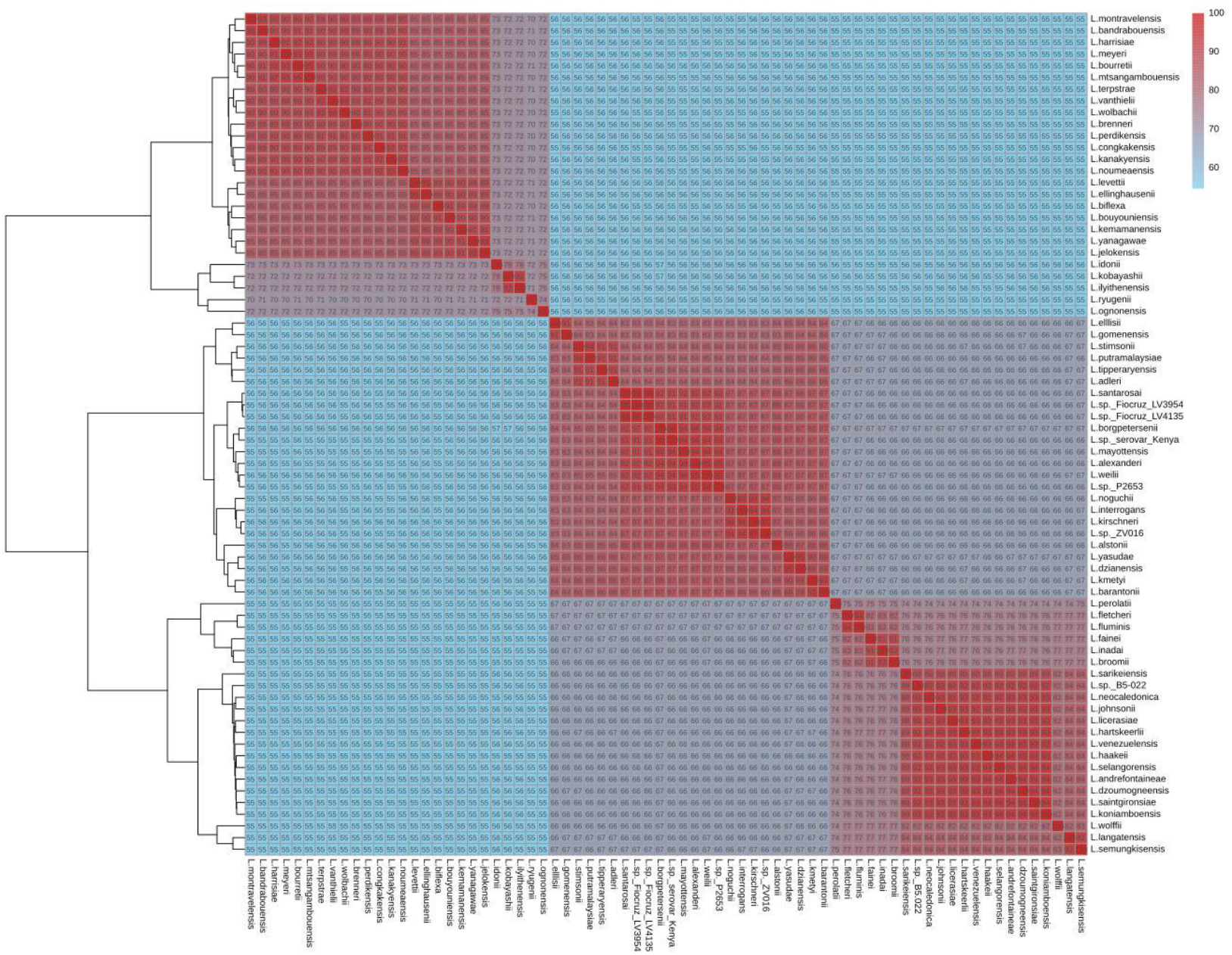
Clustering of Leptospira species on the basis of Average Amino Acid Identity (AAI) of proteins constituting core genome. Heatmap shows level of similarity on the scale of 0 to 100 where 0% similarity depicted in (light blue) and 100% similarity depicted in (red).

### 3.6 Phylogenomic analysis of *Leptospira*

Core genome of *Leptospira* genus comprised of 1038 genes of which 946 genes are single copy and rest 73 genes are multi-copy orthologs. All single copy orthologs were concatenated and analysed to establish the phylogenetic relationship among *Leptospira* species as described previously(Azam et al., 2016; Parthasarathy et al., 2017). When *Leptonima ilinii* was included as outgroup for the phylogenetic tree, single copy orthologs across all the species decreased to 705 genes. However, branching pattern in both phylogenetic trees remained same (**Supplementary Figure S5 and Figure 3**). The clustering of species is identical to the dendrogram obtained using AAI. The three phylogenetic group (**Figure 3)** cloured as red, blue and green corresponding to pathogenic, intermediate and non-pathogenic clades of AAI respectively. Each major branch of tree was supported with very high bootstrap value (100). Phylogenetic grouping of species differs with grouping of species using ANIb value. Like AAI clustering, the 5 species *Leptospira ryugenii*, *Leptospira ognonensis*, *Leptospira idonii*, *Leptospira ilyithenensis* and *Leptospira kobayashii* grouped with non-pathogenic species rather than intermediate species as in ANIb hierarchical clustering.

**Figure 3:**
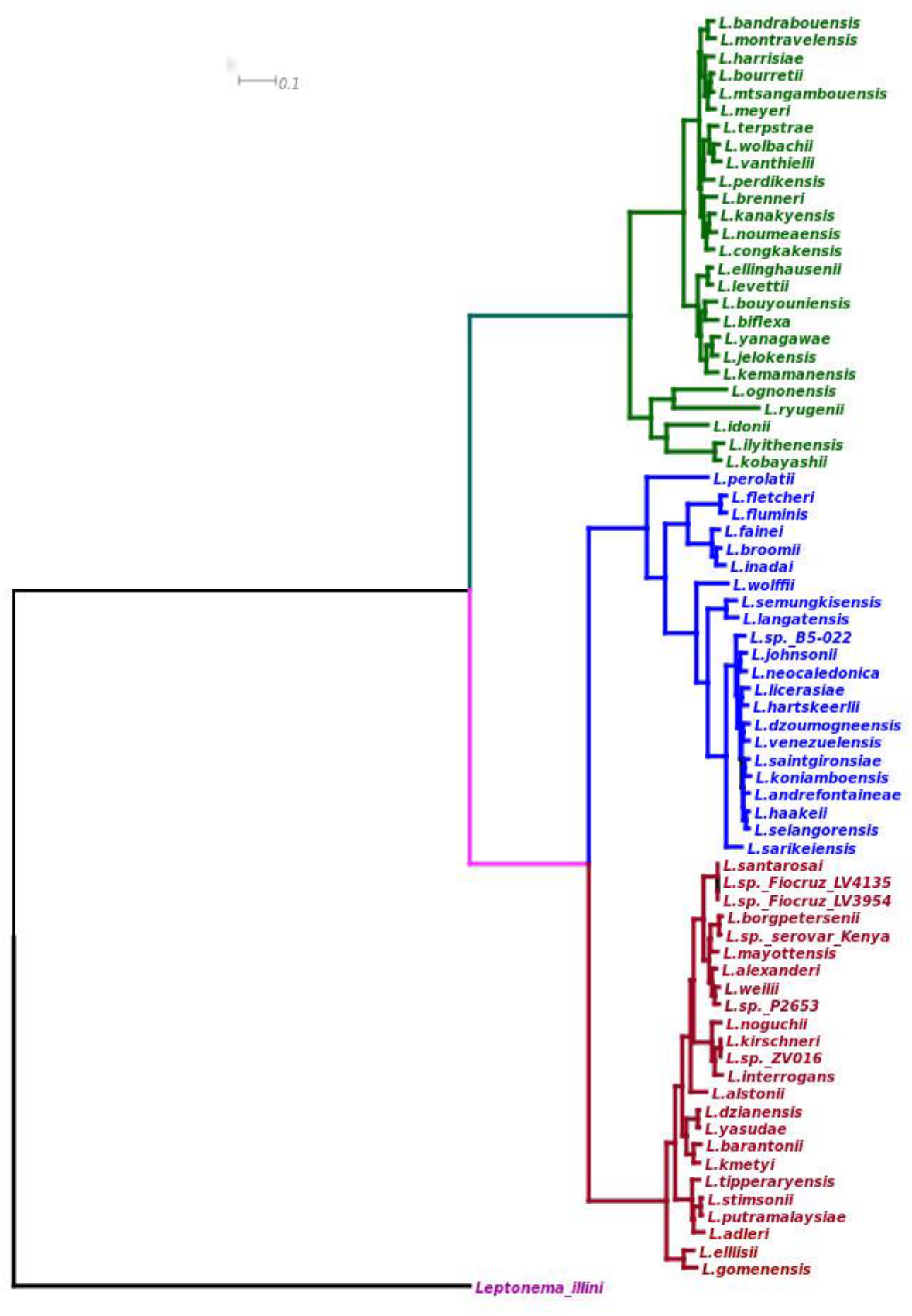
Maximum likelihood of phylogeny based on sequence similarity for the genus Leptospira on the core genome alignment. The tree is rooted with *Leptonema illini*. Phylogenomic branches are highlighted with red, blue and green colour representing with pathogenic, intermediate and saprophytic phylogenomic species respectively. Each branch is supported with 100 bootstrap values.

### 3.7 Clad-wise assessment of core and pan genome

Species grouped in each clad of phylogenetic tree were separately analysed for core, soft-core and pan genome (**Table 1**). Largest pan-genome was of pathogenic clad (12416 genes) whereas largest core and soft-core genome was of Intermediate clad. Pathogenic clad has significantly smaller core genome (11.9%) and soft-core genome 19.3% in comparison to other clads. Comparatively, cloud genes contributed higher proportion (56.9%) in pan-genome of pathogenic clad.

**Table 1.**
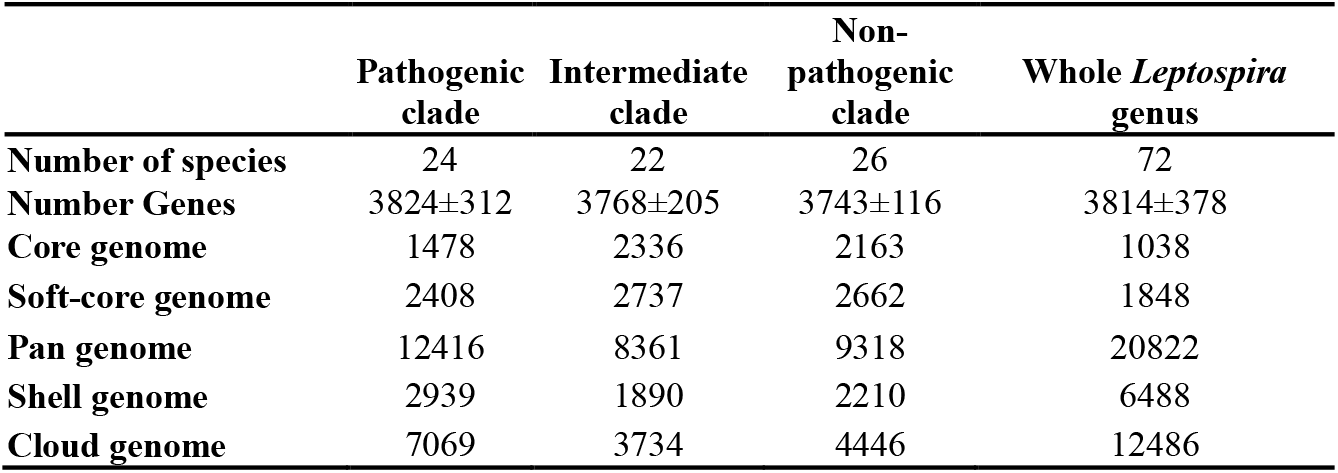
Core and pan-genome of *Leptospira* species.

|  | Pathogenic<br>clade | Intermediate<br>clade | Non-pathogenic<br>clade | Whole *Leptospira*<br>genus |
| --- | --- | --- | --- | --- |
| **Number of species** | 24 | 22 | 26 | 72 |
| **Number Genes** | 3824±312 | 3768±205 | 3743±116 | 3814±378 |
| **Core genome** | 1478 | 2336 | 2163 | 1038 |
| **Soft-core genome** | 2408 | 2737 | 2662 | 1848 |
| **Pan genome** | 12416 | 8361 | 9318 | 20822 |
| **Shell genome** | 2939 | 1890 | 2210 | 6488 |
| **Cloud genome** | 7069 | 3734 | 4446 | 12486 |

### 3.8 Functional categorization of genes in COGs

Cluster of Orthologous genes (COGs) analysis provided function of orthologous genes present in *Leptospira*. A total of 3744 (17.9%) genes of pan-genome and 721 (69.4%) genes of core genome of *Leptospira* were assigned with COGs. Clad based analysis revealed that pan-genome of non-pathogenic clad were having maximum number (2865) of COGs followed by pan-genome of Intermediate and pathogenic clade **(Supplementary Table S4 and S5)**. Functional categorization revealed (**Figure 4 a&b**) that J (translation, ribosomal structure and biogenesis), E (amino acid transport and metabolism),R (general function),T (signal transduction mechanism), M (cell wall/membrane/envelope biosynthesis), O (post translational modification, protein turnover and chaperones), C (energy production and conservation), H (coenzyme transport and metabolism) and I (lipid transport and metabolism) categories genes are generally abundant in core genomes of *Leptospira*. J category genes seems to be most abundant in core and softcore genomes of each clad whereas K (Transcription) and S (Function Unknown) categories are enriched in the pan-genome of each clad. T (signal transduction mechanism) category are enriched in pan-genome in comparison to core genome of *Leptospira*. X(mobilome), B (chromatin structure and dynamics), W (extracellular structures), Z (cytoskeleton) categories genes are sparsely present in *Leptospira*. Infact, X (mobilome) category COGs are absent in all the core genome whereas it is present in few numbers in pan-genome of each clade except pathogen clad where 65 COGs were assigned to this category.

**Figure 4:**
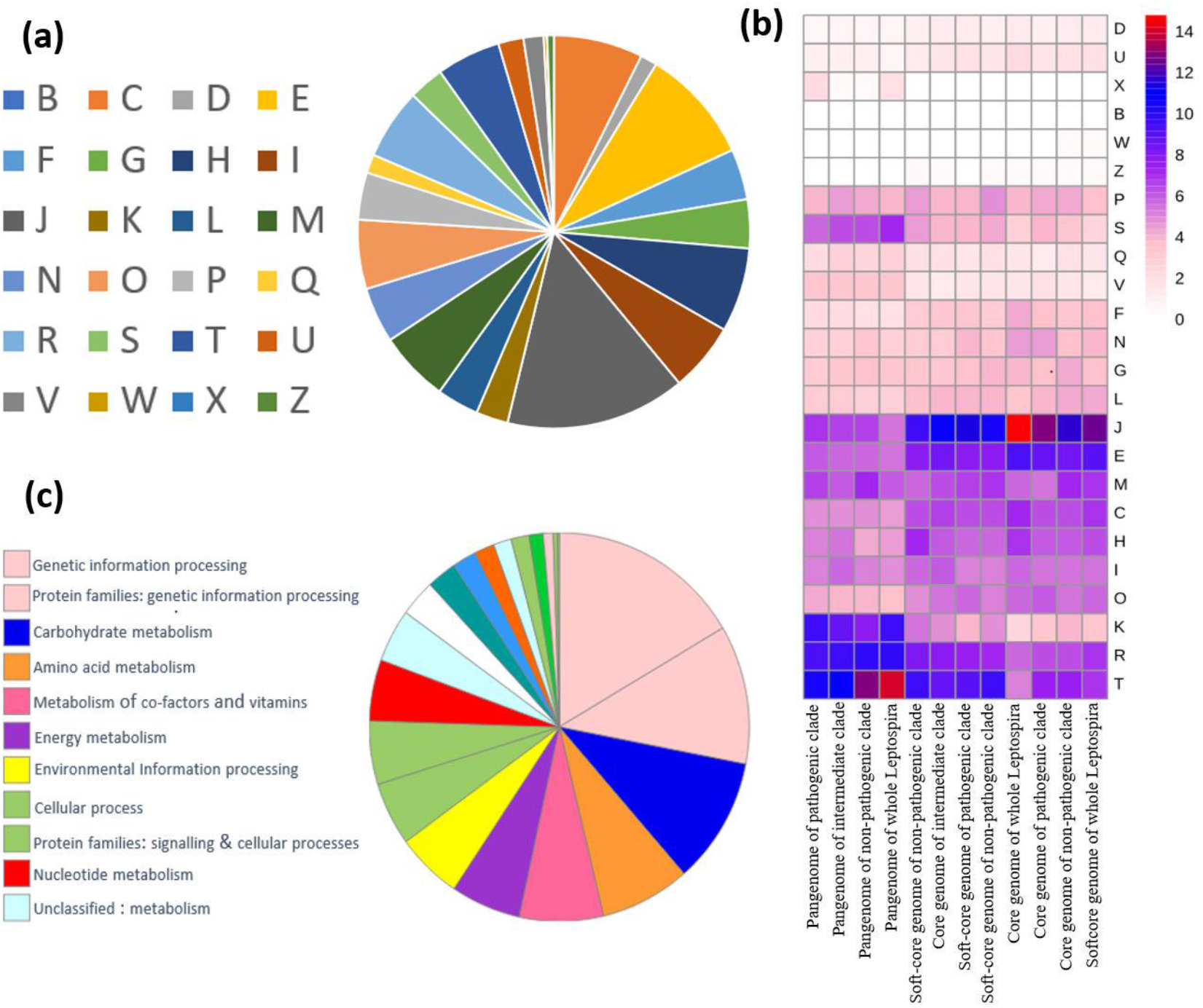
Functional classification of Leptospira genes. **(a)**: Functional categorization of genes in COGs. Cog categories are as follows: J (translation, ribosomal structure and biogenesis), E (amino acid transport and metabolism),R (general function),T (signal transduction mechanism), M (cell wall/membrane/envelope biosynthesis), O (post translational modification, protein turnover and chaperones), C (energy production and conservation), H (coenzyme transport and metabolism), I(lipid transport and metabolism), X (mobilome), B (chromatin structure and dynamics), W (extracellular structures), Z (cytoskeleton), K (Transcription) and S (Function Unknown), L (replication, recombination and repair), D (Cell cycle control, cell division, chromosome partitioning), U (Intracellular trafficking, secretion, and vesicular transport), N (Cell motility), V (Defense Mechanism), G (Carbohydrate transport and metabolism), F (Nucleotide transport and metabolism), P (Inorganic ion transport and metabolism), Q (Secondary metabolites biosynthesis, transport and catabolism). **(b)** Heatmap showing enrichment of COGs categories in different datasets. White colour represents zero or scarce presence whereas blue to red colour represents highly enriched categories **(c)** Proportion of genes assigned to different GhostKOALA functional categories.

### 3.9 *Leptospira* species are rich in metabolic pathway

Functional categorization of pan-genome using GHOSTKOALA (https://www.kegg.jp/ghostkoala/) provided landscape of known and hypothetical genes across *Leptospira* species(Kanehisa et al., 2016). Taxonomic categorization provided that 42% genes of pan-genome and almost all genes of core genome mapped (99.7%) to spirochetes category. Only 13.5% of pan-genome genes were functionally categorised (**Figure 4c**) in which most of the genes were categorized in genetic information and cell signalling and cellular processes. Like pan-genome, most of the genes of core genome were functionally categorized in genetic information processing. However, second most enriched category was “carbohydrate metabolism” followed by amino acid metabolism and metabolism of co-factors and vitamins (**Table 2**). An increase of 2.4-fold of core to pan genome was observed in pathway reconstruction in which maximum increase of 7.15 fold was observed in “organismal systems” class and lowest increase was observed in “genetic information processing” class.

**Table 2.** Reconstruction of core, soft-core and pan-genome of genus *Leptospira* and its clade with BRITE and PATHWAY algorithm of GhostKOALA.

| GhostKOALA Results | Whole *Leptospira* |  |  | Pathogenic clade |  |  | Intermediate clade |  |  | Non-pathogenic clade |  |  |
| --- | --- | --- | --- | --- | --- | --- | --- | --- | --- | --- | --- | --- |
|  | Core | Soft-core | Pan | Core | Soft-core | Pan | Core | Soft-core | Pan | Core | Soft-core | Pan |
| **Pathway Reconstruction Result** |  |  |  |  |  |  |  |  |  |  |  |  |
| Metabolism | 261 | 474 | 870 | 285 | 416 | 680 | 427 | 474 | 645 | 416 | 464 | 681 |
| Genetic Information Processing | 97 | 147 | 274 | 110 | 159 | 236 | 157 | 165 | 188 | 157 | 167 | 204 |
| Environmental Information Processing | 53 | 61 | 315 | 65 | 101 | 202 | 97 | 114 | 204 | 90 | 112 | 235 |
| Cellular Processes | 31 | 46 | 311 | 71 | 111 | 196 | 102 | 112 | 184 | 103 | 118 | 229 |
| Organismal Systems | 20 | 50 | 149 | 29 | 40 | 97 | 44 | 50 | 77 | 48 | 52 | 103 |
| Human Diseases | 26 | 43 | 159 | 38 | 49 | 103 | 48 | 53 | 93 | 51 | 59 | 110 |
| Total | 488 | 821 | 2078 | 598 | 876 | 1514 | 875 | 968 | 1391 | 865 | 972 | 1562 |
| **Brite Reconstruction Result** |  |  |  |  |  |  |  |  |  |  |  |  |
| Orthologs and modules | 587 | 967 | 2802 | 694 | 1063 | 2021 | 1064 | 1188 | 1843 | 1052 | 1200 | 2034 |
| Protein families: metabolism | 357 | 599 | 1539 | 424 | 656 | 1155 | 658 | 730 | 1061 | 653 | 726 | 1161 |
| Protein families: genetic information processing | 184 | 302 | 666 | 214 | 317 | 516 | 318 | 335 | 453 | 317 | 345 | 478 |
| Protein families: signalling and cellular processes | 123 | 204 | 742 | 151 | 229 | 502 | 209 | 242 | 469 | 215 | 251 | 532 |
| Total | 1251 | 2072 | 5749 | 1483 | 2265 | 4194 | 2249 | 2495 | 3826 | 2237 | 2522 | 4205 |

KEGG modules defined in KEGG database shows the complete functional unit of pathways. The reconstruction according to modules displays presence of pathways in genome of *Leptospira*. Core genome of *Leptospira* contains few numbers of such complete modules. However, softcore genome contains almost all short pathways either as complete or missing with one or two enzymes **(Supplementary Table S6)**. However, Pathogenic clad contain very few modules in its core genome. *Leptospira* core genome contains signature modules of Sulfate-Sulfur assimilation (module Id: M00616). Moreover, Core genome of non-pathogenic clad and intermediate clad were having genes related with signature modules other than Sulfate-Sulfur assimilation such as Nitrate assimilation (module Id: M00615), copper efflux regulator part efflux pump used in multidrug resistant, etc.

### 3.10 Presence of virulence factor in *Leptospira*

The three clad identified by AAI and phylogenetic study has been assigned to pathogenic, intermediate and non-pathogenic on the basis of virulence of *Leptopsira* species. A set of 78 *Leptospira* genes were orthologous to VFDB and can be considered as virulence factors. It was observed that all clades have virulence factor **(Figure 5a).** Core genome of pathogenic, intermediate and non-pathogenic clad having 22, 31 and 30 genes respectively. Out these, 16 genes were common and 10 genes were shared by two clads and 9 genes were unique to clad. Distribution of virulent genes in soft core was similar in numbers. Pan-genome of non-pathogenic clad were having highest number of genes (60) whereas pan-genome of pathogenic consist of only 49 genes. Functional assignment of 78 virulence genes showed that 74 genes contain 55 COGs which grouped in 18 COG categories. Maximum number of genes (32) belongs to M (cell wall/membrane/envelope biosynthesis) category. Other enriched categories were N (cell motility) and O (PTM, protein turnover and chaperons). Further, search with Victor database provided that 172, 185 and 182 virulent genes in pangenome of pathogenic, intermediate and non-pathogenic clade respectively (**Figure 5b**). 198 genes were identified when searched in pangenome of Leptospira. Parallelly, list of 304 non-redundant genes (**Figure 5c, Supplementary Table S7**) considered as virulent or has role in host-pathogen interactions was compiled manually from published literatures. When all datasets (VFDB, Victor and known genes) were compared, a total of 496 orthologs involved in virulence or host pathogen interactions, were obtained. 12 genes were common in all datasets and 36 genes were common between Victors and VFDB database.

**Figure 5:**
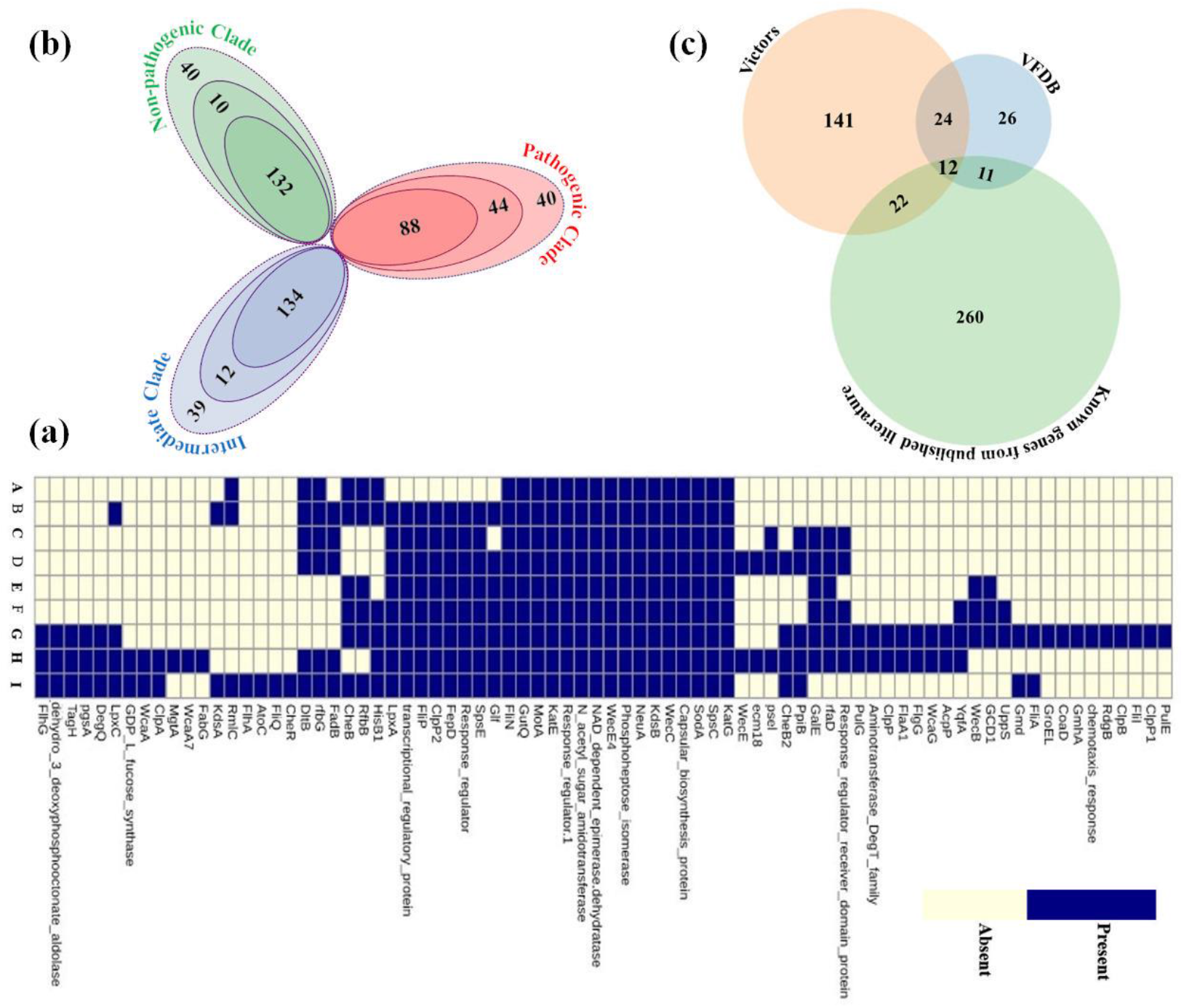
virulence gene in leptospira: **(a)** virulent genes identified using VFDB**. Distribution of genes in different dataset has been presented as heatmap.** Datasets are A-core genome of pathogenic clade, B-softcore genome of pathogenic clade, C-core genome of intermediate clade, D-softcore genome of intermediate clade, E-core genome of non-pathogenic clade, F-softcore genome of non-pathogenic clade, G-pangenome of non-pathogenic clade, H-pangenome of intermediate clade, I-pangenome of pathogenic clade. Blue represent presence and light yellow represent absence of virulent genes in dataset. **(b)** Virulent genes identified using Victors database in each clade. Virulent genes identified in core genome (inner ellipse), softcore genome (middle ellipse) and pangenome (outermost ellipse) are represented in each petal. **(c)** Venn diagram showing intersections of sets of virulent genes identified in Leptospira using VFDB, Victors database and known gene curated from published literature.

### 3.11 Identification of clad specific genes

Clad specific genes were identified by comparing core genome of each clade with the pan-genome of other clades **(Supplementary Table S8)**. Twenty-nine genes were found to be specific to pathogenic clad whereas 253 genes were specific in non-pathogenic clad. Intermediate clad were having 43 such specific genes. Moreover, 8 genes were exclusively absent in pathogenic clad but present in all intermediate and non-pathogenic species. On the other hand, 110 genes were found in all intermediate and pathogenic species and absent in saprophytic species **(Figure 6)**. Further these clad specific genes were searched for their KOs in KEGG database. Pathogenic specific genes (29) were not assigned with any KOs, however only five of intermediate specific genes (43) were assigned KOs which was associated with metabolism. Similarity search with known virulence factor database (VFDB) revealed that none of the clad specific genes belongs to virulence factor.

**Figure 6:**
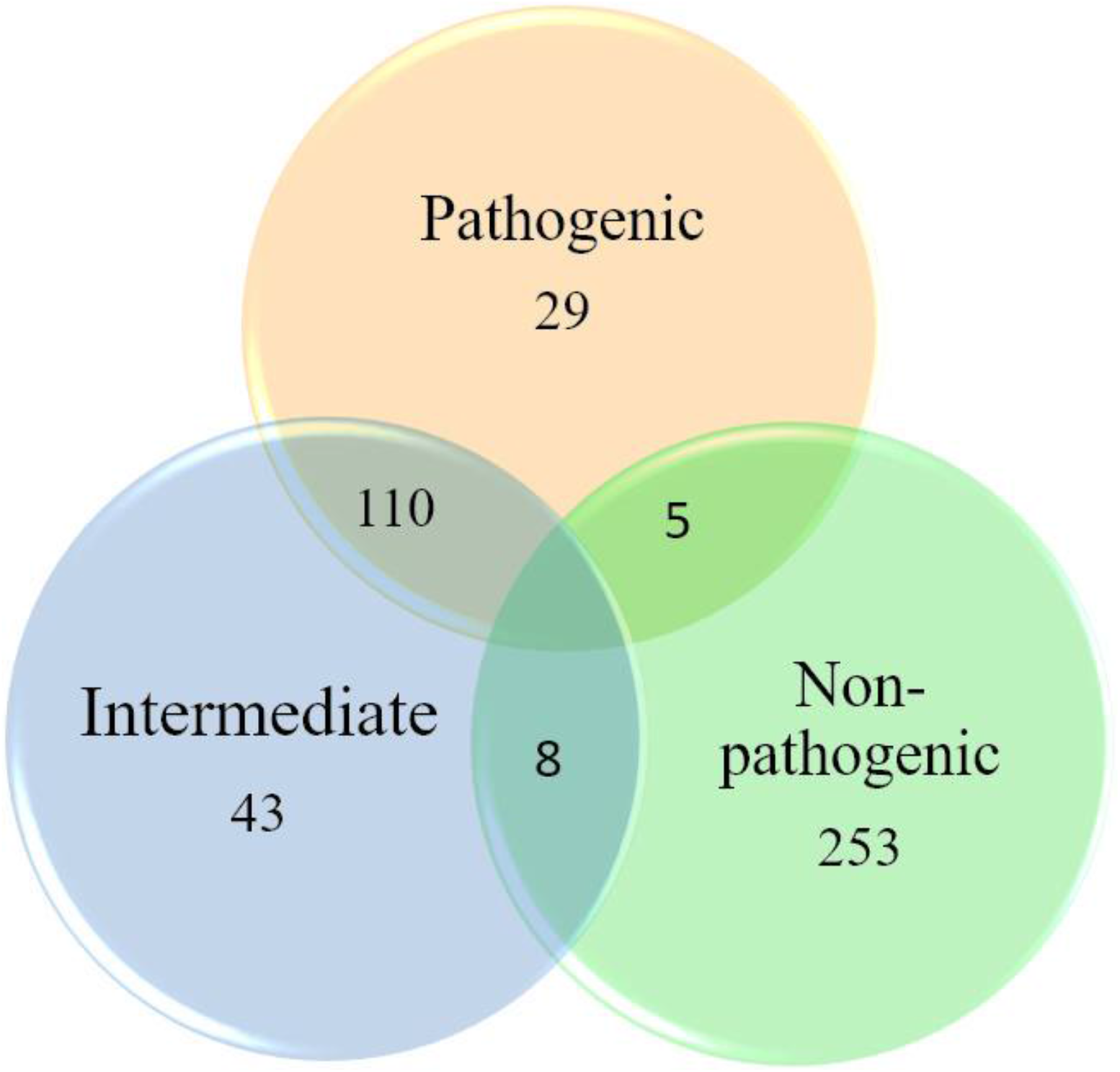
Venn diagram showing specific genes exclusively present in different clade of *Leptospira*.

## 4. Discussion

NCBI RefSeq is a curated database which comprehensively analysed the genome using **Prokaryotic Genome Annotation Pipeline (**PGAP). This provides a stable reference genome for annotation, gene identification and characterization of each species. However, name and taxonomic position of species are provided by the sequence submitter which is generally determined using 16S RNA or MLST analysis. After WGS, various organisms are redefined and new taxonomic position has been assigned. For example bacteria of *Shewanella Taxonomy*(*Thorell et al., 2019*) such as *S. arctica*(*Kim et al., 2012*) earlier defined as *S. frigidimarina* (*Bozal et al., 2002*). Normally a polyphasic approach has been used to assign the evolutionary relationship and classification on the basis of phenotype, genotype and chemotaxis parameter. Assignment of new isolate to species is based on the genotypic parameter of DNA-DNA hybridization >70%. In case DNA-DNA hybridization is <70%, species are assigned to genus on various serological and chemotaxis parameter. When DDH is translated at whole genome similarity level, it was established that average nucleotide identity (ANI) of >95% indicates that genomes belong to the same species(Jain et al., 2018). Although, ANI was not defined for genus delineation, however, it was observed that minimum of 67% of ANI values in *Chlamydiales* (*Pannekoek et al., 2016*) and 68% ANI value in the *Alteromonadales* was found between a pair of species of same genera(Barco et al., 2020; Qin et al., 2014). In our study, well annotated reference genome of 73 species of *Leptospira* was downloaded from RefSeq database. *Leptospira macculloughii* was outlier as expected because it was sequenced from mixed culture of *L. meyeri* and *L. levetti*(Thibeaux et al., 2018), hence, excluded from analysis. Average Nucleotide Identity(ANI) between pair of species fall in range of 68-95% which is very much in concordance with known ANI parameter except 8 pairs which shows > 95% identity. Minimum observed value of ANI between species suggest that all genomes belongs to genus *Leptospira*. Similar results were observed for genera of *Chlamydiales* (*Pannekoek et al., 2016*) and *Alteromonadales*(*Qin et al., 2014*). Out of the 8 pairs showing > 95% ANI values, three pairs i.e. *L. putramalaysiae* vs *L. stimsonii*, *L. weilli* vs L. sp. P2653 and *L. yasudae* vs *L. dzianensis* showed less than 96% ANI and <70% DDH in GGDC analysis and can be considered as separate species. In fact, kim et al. suggested that interspecies ANI value can go upto 96%(Kim et al., 2014). L. sp. Fiocruz LV4135, L. sp. Fiocruz LV3954, and L. santarosai showed more than 98% nucleotide identity and also >70% DDH establish that these are strains of the same species. Similarly, L. sp. Serovar Kenya vs *L. borgpetersenii*, *L. kirschneri* vs L. sp ZV016 are pair of strains and belongs to same species. These strains which are represented as species in the study are recently sequenced. Many strains of *L.santarosai*, *L.weilii and L. interrogans* was misclassified earlier as well(Vincent et al., 2019). Hence, our study indicates that only 68 species of *Leptospira* are actually sequenced at the time this study.

*Leptospira* genus consist of diverse species and exist in heterogeneous environment. Such genus have large and open pan-genome. Generally, closed pan genomes are achieved in species which occur in few habitat and have low capacity to acquire genes(Rouli et al., 2014). Pan-genome of *Leptospira* contains 20822 genes which is similar to the pan-genome of other genus having diverse species such as *lactobacillus*(Inglin et al., 2018). Earlier study has reported pan genome on species level which are generally of 7000-8000 genes (Kurilung et al., 2019). It is obvious that a genus will have larger pan-genome than its species. As far as core genome is concerned, diverse genus such as *Streptococcus* (*Gao et al., 2014*), *Helicobacter pylori*(*Lu et al., 2014*) have comparatively small core genome. However, core genome assessed in the study is relatively larger but in accordance with previous study which reported core genome of Leptospira contain 1023 genes(Xu et al., 2016). Hierarchical clustering of species on basis of ANI and AAI reveal different clustering pattern. ANI clustering is based on pan genome and revealed 4 clusters whereas clustering on AAI value of core and pan-genome shows 3 optimum clusters **(Figure 2).** Optimal clustering is supported by silhouette width calculations as well. Further, phylogenomic analysis also provide tree consistent with core and pan genome based dendogram on AAI values. The major difference in the clustering pattern was that intermediate cluster on AAI values corresponds to two groups i.e. C and D of ANI dendrogram except five species of group D. These five species of group D are *Leptospira ryugenii*, *Leptospira ognonensis*, *Leptospira idonii*, *Leptospira ilyithenensis* and *Leptospira kobayashii* which grouped to non-pathogenic cluster in AAI based dendogram and phylogenetic tree. These five species are proven to be saprophytic in nature(Masuzawa et al., 2019; Masuzawa et al., 2018; Saito et al., 2013; Vincent et al., 2019) and should be grouped with saprophytic species while clustering. Thus, core and pan genome based clusters on AAI value and phylogenetic analysis, grouping these five species with other non-pathogenic species shows true grouping. The AAI based clustering pattern is also supported with previous study by **Vincent, et al**.^20^, where these five species are grouped in saprophytic clad. In contrast to ANI based clustering, all intermediate species are expected to be grouped together as in AAI based dendogram and phylogenetic tree. All intermediate species has been grouped into one clad in previous study as well(Vincent et al., 2019). This indicate that grouping of species on pangenome based ANI value provide erroneous clusters. In fact, Vincent et al. proposed two clades P (pathogenic) and S (saprophyte) consisting 4 sub clades i.e. P1 (pathogenic), P2 (pathogenic), S1 (saprophyte) and S2 (saprophyte) for Leptopsira genus(Vincent et al., 2019). Similar phylogenetic pattern were observed on AAI based dendrograms or phylogenetic tree in our study where P1 corresponds to pathogenic, P2 corresponds to intermediate and S1 & S2 merged to non-pathogenic (saprophyte) clusters. Previous study also calculated for core and pan-genome for each clade (P1, P2, S1 and S2) separately. P1, P2, S1 and S2 were having core genomes of 1560, 2305, 2564, 2385 clusters and pan-genome of 11224, 8010, 7718 and 5939 clusters respectively(Vincent et al., 2019). Clade-wise distribution of genes showed pathogenic clade has least number of core genes. In fact, core genome of each clad were shown nearly closed. Similar results were observed in our study as well. In pathogenic clad which correspond to P1, hardly 70 genes were decreased even after the addition of 7 species. It is normal that core size decreases with addition of genomes until it become completely closed(Inglin et al., 2018; Rouli et al., 2015). Similarly, intermediate clad shown comparable number of core clusters with P2 clad. But non-pathogenic core cluster was much lesser because it represent the merged clad of both saprophyte (S1 and S2). On the other hand, total cluster increases by more than 1200 in the pan genome of pathogenic clad. This huge increase in cluster was contributed by 7 genomes and shows that pan-genome is still open. S1 and S2 clades were having total of 26 species which are the same constituting non-pathogenic clade in this study. After merging the clads, pan-genome were reduced to 9318 clusters in our study. This huge reduction of cluster from combined pan-genome (7718+5399) shows that both S1 and S2 shared a very high number of common genes. Apart from genes in core and pan-genome, cloud genes show the presence of unique gene which are generally present in one or two species of genus. These genes may provide the unique features to a species(Bochner, 2009). High number of such cloud genes in Pathogenic clad infer that many of the genes has been acquired through horizontal gene transfer(Sarkar and Guttman, 2004).

Functional categorization of genes using COG database are on the basis of conserved orthologs. Core genomes are generally conserved and rich of essential genes(Merkl, 2006), (Liu et al., 2012). In comparison of pangenome, high proportion of core genes in this study were functionally characterized and assigned to COGs category. Similar results have been observed in previous studies on Staphylococcus aureus (Zubair et al., 2015) *Helicobacter*^43^. J category enriched in core genome across the *Leptospira* clades support previous studies that translational machinery remains conserved and close to original form in evolution(Liu et al., 2012; Ouzounis and Kyrpides, 1996). In contrast, X category COGs contains mobilome related genes, were found only in pan-genome of pathogenic clade. These genes can be attributed to phenotype of the clade as genes in the mobilome category were overwhelmingly predicted as Horizontally transferred genes and has role in pathogenicity(Nakamura, 2018). Further, KEGG pathway annotation also revealed that most of the genes of core genome are related with genetic information and processing. Various signature module of KEGG database like Sulfate-Sulfur assimilation, Nitrate assimilation etc. observed in the study has also been reported for *Leptospira* (Valdes et al., 2003; Zhang et al., 2018). In contrast, most of clade specific genes can’t be categorized for any functional class. These lineage specific genes are subset of core genes of their respective clades. Basically, these are also the gene gain in particular clade (Yu, 2009). Study has shown the use of such clade specific or lineage-specific genes as marker and their role in virulence and niche adaptation(Gori et al., 2019). In fact, clade specific genes indicate recent directional evolutionary pressure on the genome as well(Wolfe et al., 2015). Clade specific genes extracted in our study can be used as marker for metagenomic community and may help to assign new *Leptospira* species to specific clade. Howeve, such role of lineage specific genes in *Leptospira* needs to be validated in wet-lab.

*Leptospira* are known for their virulence in wide range of host. This was reflected in analysis when species of pathogenic clade have high number of species specific/unique genes causing a large pan-genome. On the contrary, decrease in core genome of pathogenic clade in comparison to other clade shows reduction of core genes in the species(Liu et al., 2012). Earlier also it was reported that pathogenic *Leptospira* undergo reductive evolution and lose unnecessary genes (Xu et al., 2016). Similarly, genome reduction has been observed in other pathogenic bacteria as well as symbiotic bacteria (*Segerman, 2012*),(McCutcheon and Moran, 2011).

Many pathogenic bacteria resulted high number of virulent genes when searched against virulence factor database (VFDB) such as *Pasturella multocida (Cao et al., 2017), Pseudomonas (Vodovar et al., 2006), salmonella(Holt et al., 2009), Brucella (Crasta et al., 2008)* etc. However, less number of virulent orthologs were identified in *Leptospira* when searched against VFDB. On the other hand, Surprisingly, many of the virulent genes were found in the species of non-pathogenic clade as well. In fact, some virulent genes were part of the core genome of non-pathogenic clade. This result reconfirms earlier report of virulent genes in non-pathogenic *Leptospira*(Kurilung et al., 2019). Various known virulence factors of *Leptospira* were not annotated in VFDB search analysis such as Lig proteins (Lig A, B, C) (Cerqueira et al., 2009), Lipoproteins (LipL32, LipL45, LipL71)(Oliveira et al., 2011), Len proteins (Len A, B, C, D, E, F)(Stevenson et al., 2007) and Tly proteins (Tly A, B, C)(Carvalho et al., 2009) etc. This shows that VFDB is devoid of known virulent genes of *Leptospira* genus and not a comprehensive database. Since, *Leptospira* is a zoonotic pathogen, we explored recently developed Victors database which cover virulent factors from both animal and human pathogen. Although, Victors provided high number of virulent genes (198) but it excluded many of the known virulent genes such as Lig and Len family of proteins. This indicates victor is more comprehensive than VFDB for Leptospira but alone is not sufficient. This tempted us to extensively search literature and manually prepare a list of all reported virulent factors or genes showing role in host-pathogen interaction. Large proportion of manually curated genes remained unique and was not reported by either Victors or VFDB. Hence, a non-redundant orthologs from all combinatorial approach may decipher the complete virulome of *Leptospira*.

## 5. Conclusion

Comparative genomics analysis of 72 *Leptospira* species available at RefSeq database provide insight about evolution and diversity of the genus. Clustering genus into clades provide an opportunity to dissect phenotype and adaptation to virulence of the *Leptospira* species. Clade specific genes needs to be verified in future to use as a marker or panel of markers for associating species to different clade and predicting the level of virulence. Virulent factors mined in the study will help in targeting pathogenic *Leptospira* with new tools and diagnostics. The study also emphasizes the development of new or updated database and algorithm for automated and comprehensive detection of virulent factors in *Leptospira* and other such species. The overall knowledge gained about genomic structure, phylogeny, gene functions and virulence will contribute in future research for developing new diagnostic, vaccine and drug target for *Leptospira*.

## Acknowledgements

This work was supported from DBT grant (BT/PR21430/ADV/90/246/2016) funded to SMF from Department of Biotechnology, Ministry of Science and Technology, Government of India. Financial support from NIAB core funds to SMF and SA is duly acknowledged. The authors would like to thank Director, NIAB for providing all the facilities and his constant support and encouragement.

## Author’s contributions

SA and SMF conceived the idea and designed the study. MA, MK and SA performed the analyses. RS and MSB provided critical insights and suggestions in various parts of the data analysis. SMF and SA wrote the initial draft, RS and MSB helped in editing the manuscript. All authors read and approved the final version of manuscript.

## Competing interests

The authors declare no competing financial interests.

## Supplementary Materials

***Supplementary material 1***: The document contains, Supplementary Figure S1: Distribution of genes in sequenced genomes of Leptospira and its Bubble analysis. Supplementary Figure S2: Clustering of Leptospira species on the basis of Average nucleotide Identity (ANI). Supplementary Figure S3: Optimal number of clusters (k) determined using average silhouette width. Supplementary Figure S4: Clustering of *Leptospira* species on the basis of Average Amino Acid Identity (AAI) of proteins constituting pangenome. Supplementary Figure S5: Maximum likelihood of phylogeny based on sequence similarity for the genus *Leptospira* on the core genome alignment. Supplementary Table 4: Categorization of *Leptospira* genes in COGs. Supplementary Table 6: Reconstruction according to modules for different clades in *Leptospira*. Supplementary Table 8: Clade specific genes of *Leptospira* genus.

***Supplementary Table S1:*** Details of 73 species of *Leptospira* downloaded from RefSeq database of NCBI.

***Supplementary Table S2:*** Average Nucleotide identity between 72 different species of *Leptospira*.

***Supplementary Table S3***: DDH values between 72 different *Leptospira* genomes in our study.

***Supplementary Table S5:*** Leptospira genes in different COG categories.

***Supplementary Table S7:*** List of *Leptospira* genes involved in virulence and host-pathogen interaction.

## Supplementary Information

**The supplementary file contains Figure S1, S2, S3, S4, S5 and its legend and Supplementary Table ST4, 6 and 8. ST1, 2, 3, 5 and 7 are attached as excel sheet**

**Supplementary Figure S1.**
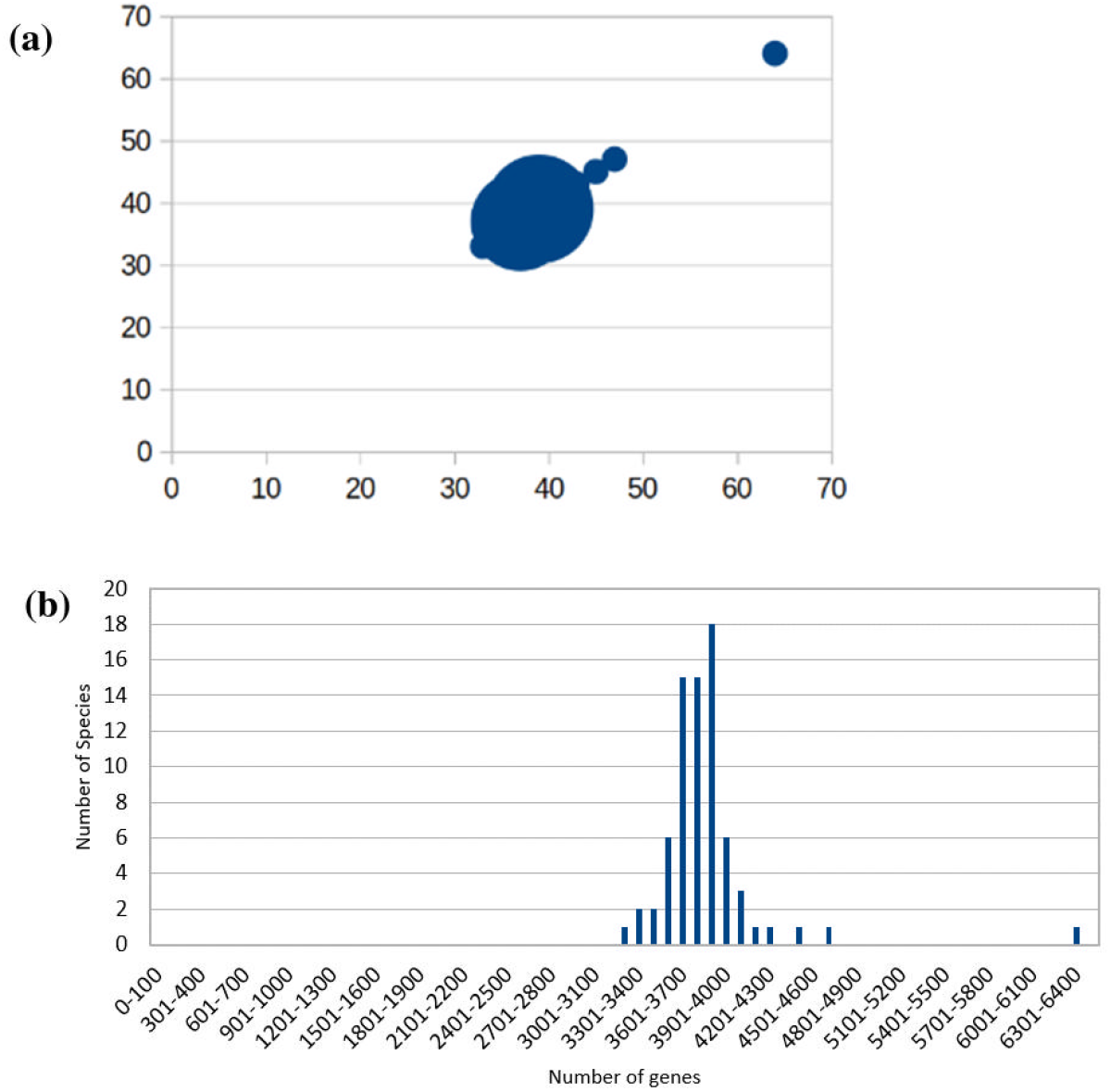
Distribution of genes in sequenced genome of Leptospira. (a) Bubble analysis shows each dot belongs to one species (b) Bar-plot showing number of species in different gene bins.

**Supplementary Figure S2:**
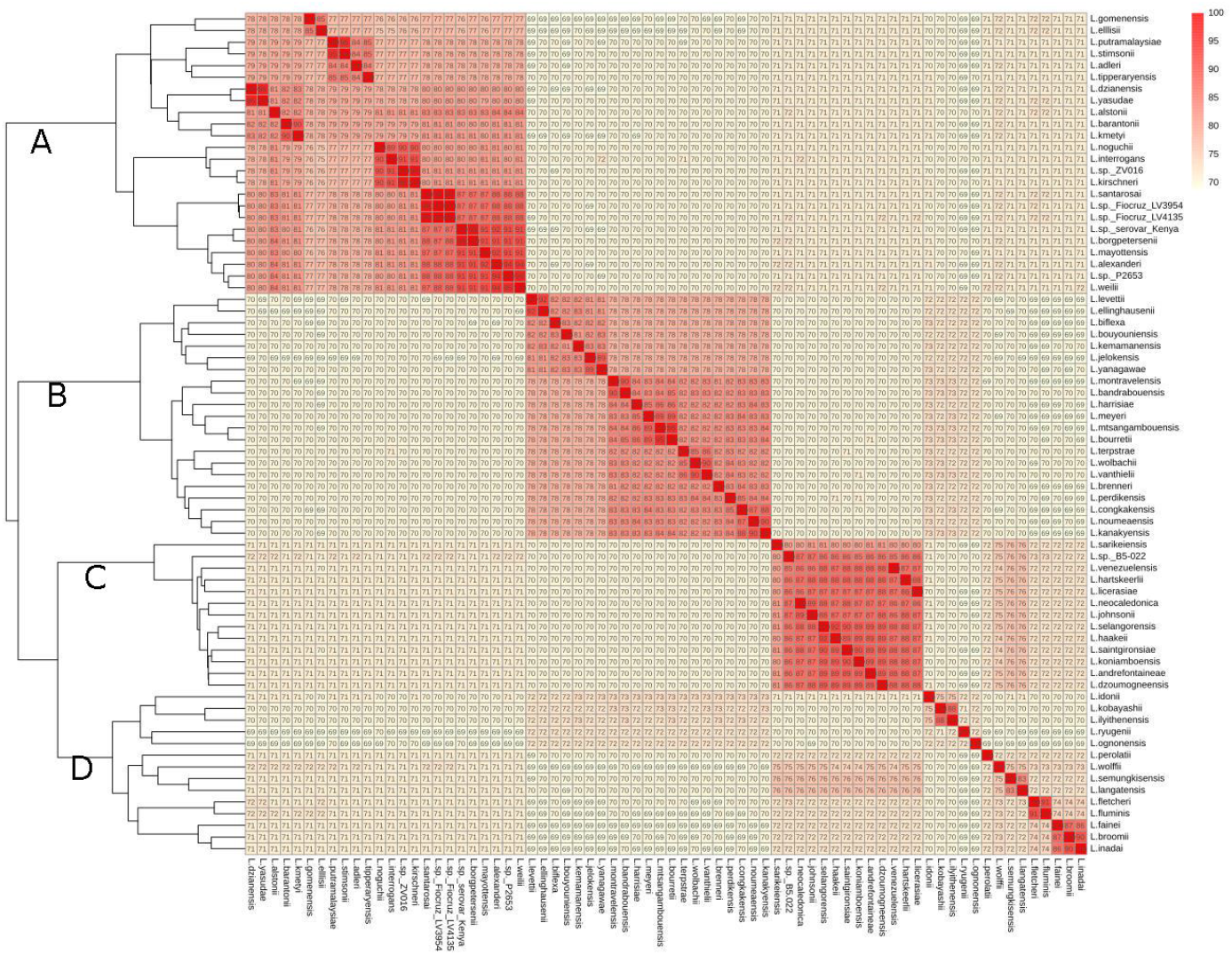
Clustering of Leptospira species on the basis of Average nucleotide Identity (ANI). Heatmap shows level of similarity on the scale of 0 to 100 where 0% similarity depicted in (yellow) and 100% similarity depicted in (red). 4 clusters observed was labelled as A, B, C and D.

**Supplementary Figure S3:**
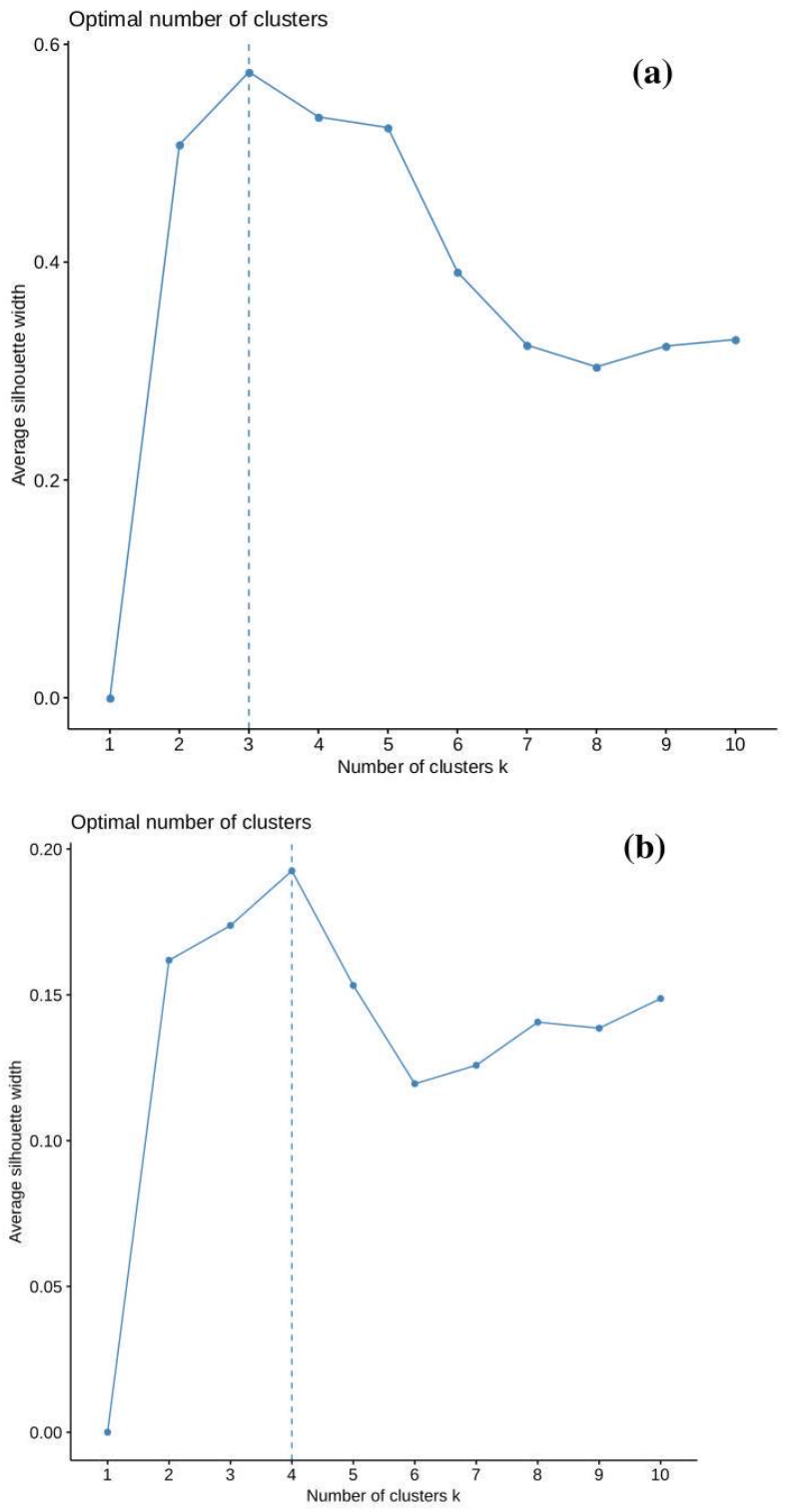
optimal number of clusters (k) determined using average silhouette width. Number of clusters observed on different average silhouette width for AAI values (a) and for ANI values (b)

**Supplementary Figure S4:**
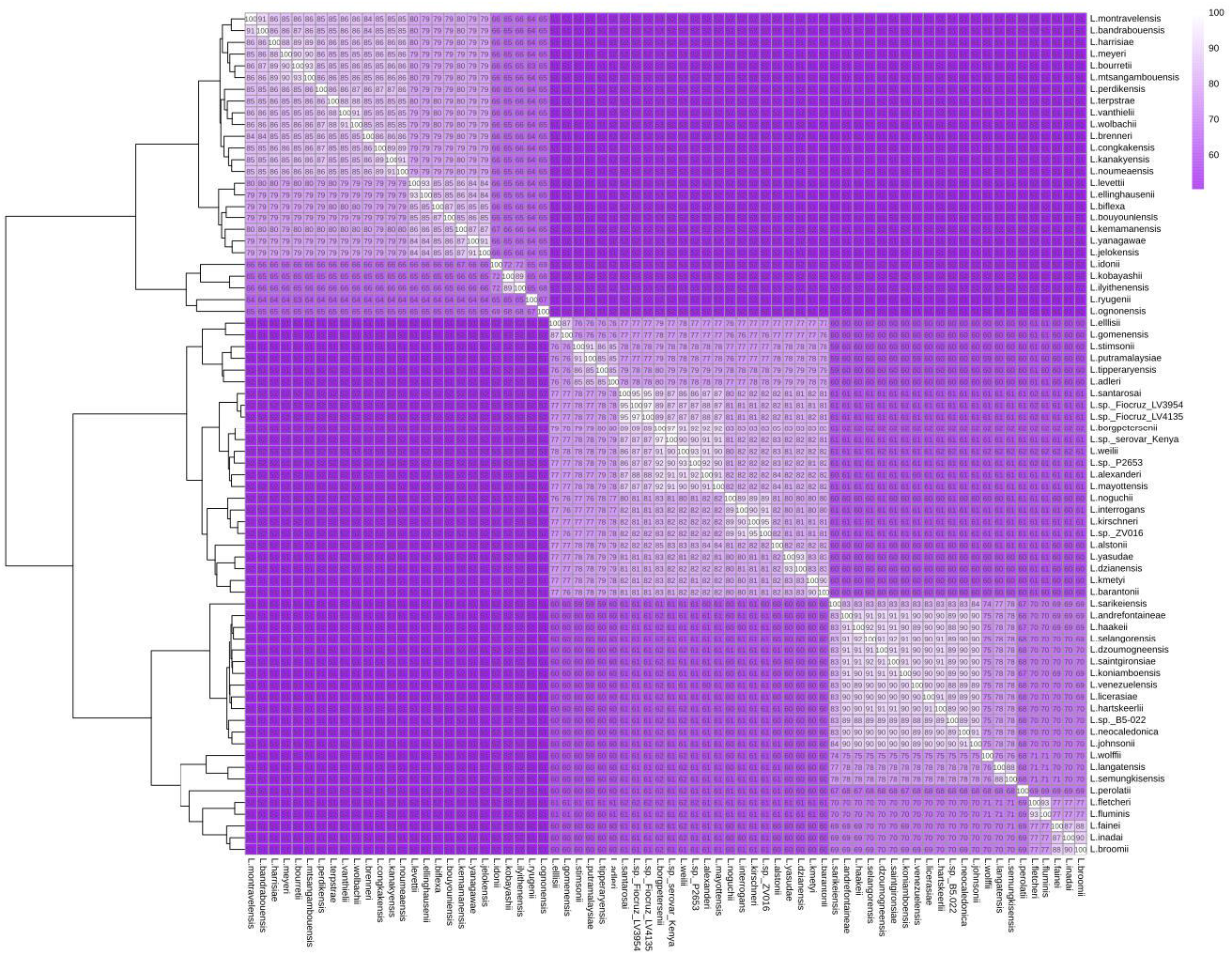
Clustering of Leptospira species on the basis of Average Amino Acid Identity (AAI) of proteins constituting pangenome. Heatmap shows level of similarity on the scale of 0 to 100 where 0% similarity depicted in (purple) and 100% similarity depicted in (white).

**Supplementary Figure S5:**
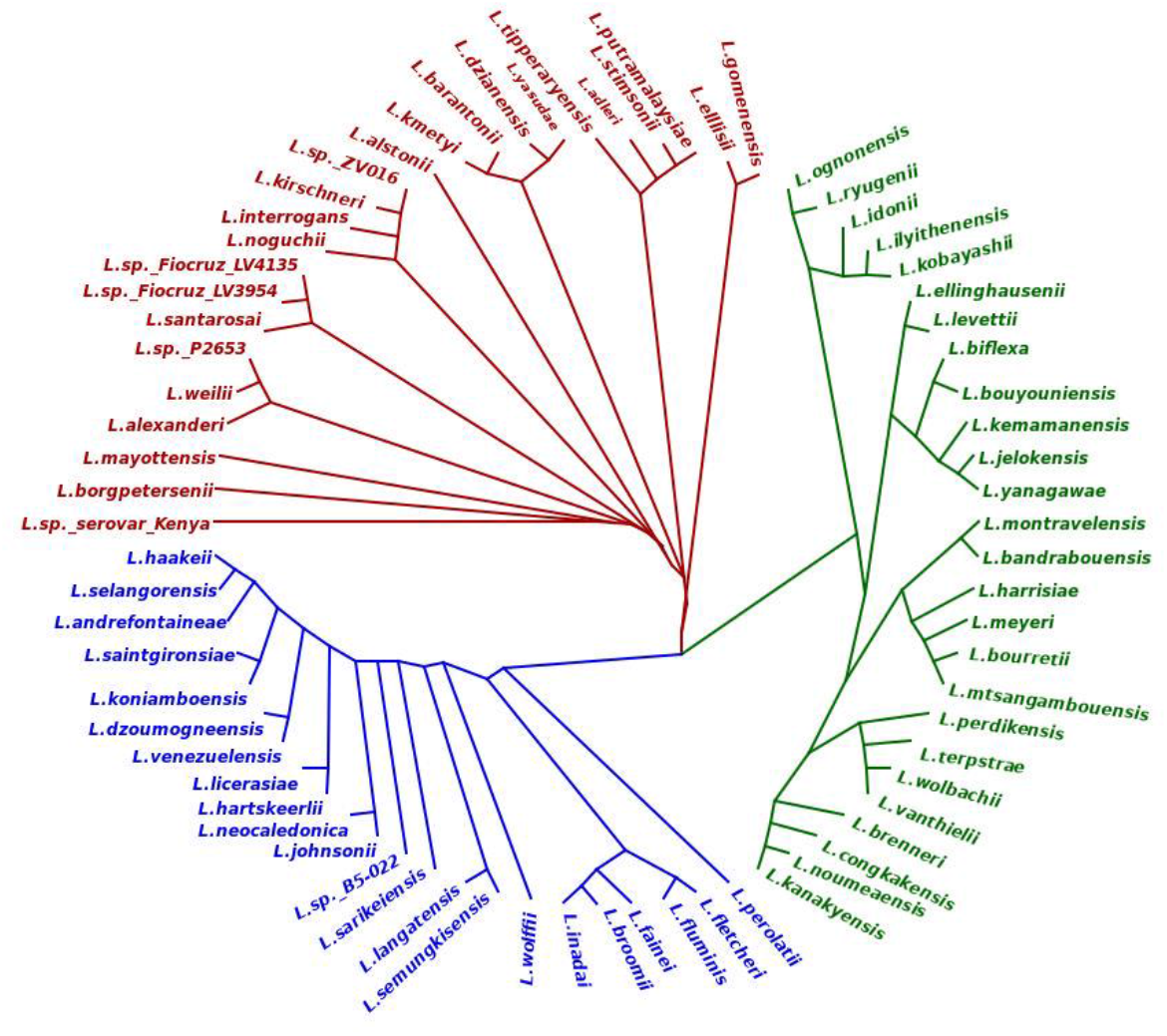
Maximum likelihood of phylogeny based on sequence similarity for the genus Leptospira on the core genome alignment. The tree is unrooted. Phylogenomic branches are highlighted with red, blue and green colour representing with pathogenic, intermediate and saprophytic phylogenomic species respectively. Each branch is supported with 100 bootstrap values.

**Supplementary Table 4.**
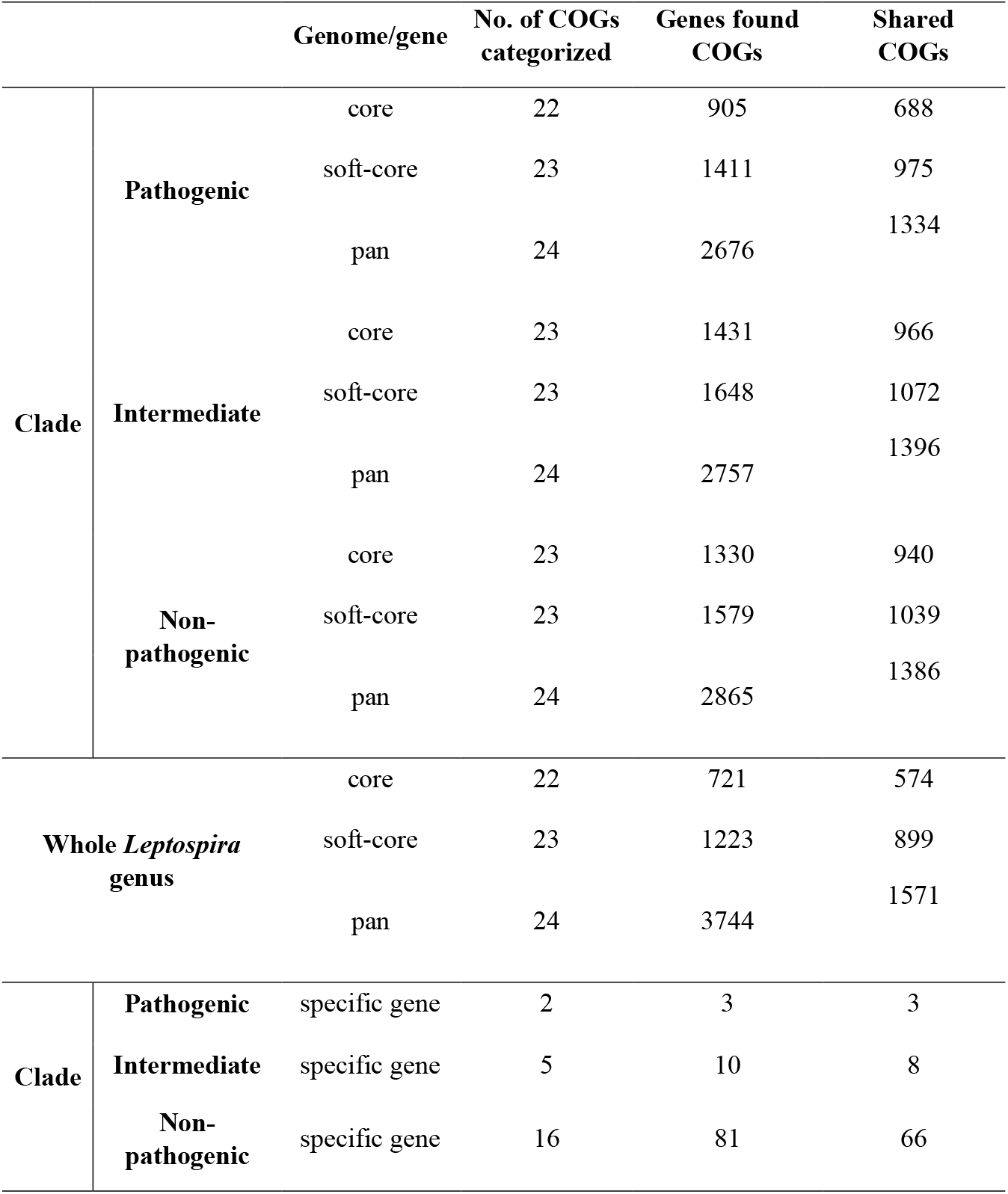
Categorization of *Leptospira* genes in COGs.

**Supplementary Table 6.** Reconstruction according to modules for different clades in *Leptospira*.

|  | Core genome |  |  | Soft-core genome |  |  | Pan genome |  |  |
| --- | --- | --- | --- | --- | --- | --- | --- | --- | --- |
|  | Complete | 1 block missing | 2 block missing | Complete | 1 block missing | 2 block missing | Complete | 1 block missing | 2 block missing |
| **Pathogenic clade** | 8 | 29 | 23 | 30 | 30 | 19 | 35 | 38 | 24 |
| **Intermediate clade** | 29 | 31 | 20 | 32 | 34 | 22 | 37 | 36 | 23 |
| **Non-pathogenic clade** | 30 | 33 | 19 | 36 | 28 | 19 | 45 | 28 | 27 |
| **Whole *Leptospira* genus** | 8 | 24 | 18 | 30 | 26 | 21 | 42 | 36 | 29 |

**Supplementary Table 8.**
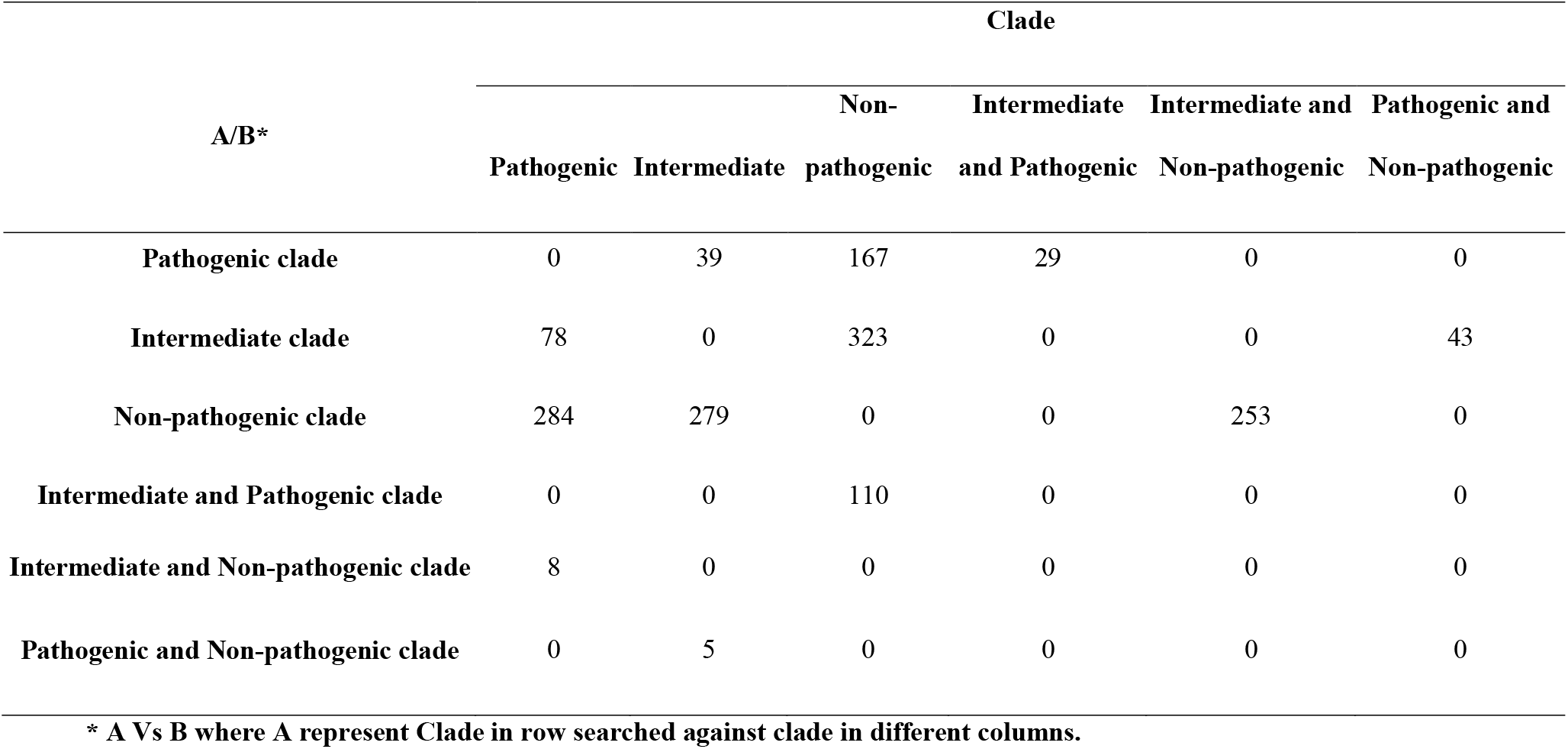
Clade specific genes of Leptospira genus.

## Notes

### Competing Interest Statement

The authors have declared no competing interest.

